# Rapid loss of genetic variation and increased inbreeding in small and isolated populations of Norwegian wild reindeer

**DOI:** 10.1101/2024.07.09.598942

**Authors:** Brage B. Hansen, Bart Peeters, Øystein Flagstad, Knut Røed, Michael D. Martin, Henrik Jensen, Hamish A. Burnett, Vanessa C. Bieker, Atle Mysterud, Xin Sun, Steeve D. Côté, Claude Robert, Christer M. Rolandsen, Olav Strand

## Abstract

Wildlife responses to habitat loss and fragmentation are a central concern in the management and conservation of biodiversity. Small and isolated populations are vulnerable, both due to demographic and genetic mechanisms, which are often linked. Thus, understanding how (changes in) genetic diversity, effective population sizes, and levels of inbreeding relate to population size and degree of isolation is key for developing effective conservation strategies. High-density Single Nucleotide Polymorphism (SNP) arrays represent an increasingly cost-efficient tool to achieve the data needed for such analysis. Here, we present the development of a novel 625k SNP array for reindeer *Rangifer tarandus* and apply this array to assess conservation genetic issues across thirteen Norwegian wild reindeer populations of varying size, isolation, and genetic origin (i.e., semi-domesticated reindeer origin or a mix of wild reindeer and semi-domesticated reindeer origins). Many of these populations are currently completely isolated, with no gene flow from other populations. We genotyped n = 510 individuals sampled by hunters and found that variation in population size across the populations largely predicted their (recent loss of) genetic variation (observed heterozygosity, H_o_), as well as effective population size (N_e_) and (change in) level of recent inbreeding. For the smallest and most isolated populations, with total population sizes of <50-100 individuals and a high and increasing level of recent inbreeding, estimated loss of genetic variation was as high as 3-10% over the time span of a generation or less, and estimated N_e_ was as low as six individuals. With the current level of isolation and associated lack of gene flow, and considering their already low genetic diversity, these populations are hardly viable – neither demographically nor genetically – in the long term. These results have direct relevance for the management of Norwegian wild reindeer, recently red-listed as ‘Near Threatened’. Yet, these genetic challenges, characterizing many of the small ‘wild reindeer’ populations in Norway, have been largely ignored by management thus far. Mitigation efforts such as reducing barriers would introduce substantial conservation dilemma due to the aim of avoiding further spread of chronic wasting disease (CWD), as well as potential further domestic introgression into populations with genetically wild reindeer (or mixed) origin. Nevertheless, our cost-efficient and high-density SNP array especially designed for reindeer and caribou offers a powerful genetic tool to include in future monitoring, providing important contributions to management and conservation decisions.

## Introduction

The conservation of genetic biodiversity is a critical challenge under accelerating habitat loss and fragmentation, as well as climate change (Sala et al., 2000; Fischer & Lindenmayer, 2007). In particular, small and isolated populations are vulnerable as both genetic and demographic factors can drive them toward extirpation (Brook et al., 2008). Genetic diversity is crucial for the adaptive potential, as it allows species to, e.g., respond to environmental change and tackle diseases (Hughes et al., 2008; Reed & Frankham, 2003). For example, low genetic diversity of cheetah (*Acinonyx jubatus*) populations makes them susceptible to disease outbreaks (O’Brien et al., 1985), and the small population size of the Florida panther (*Puma concolor coryi*) has led to significant inbreeding depression (Johnson et al., 2010). Inbreeding, the mating of close relatives, increases homozygosity, which can expose deleterious recessive alleles and reduce overall fitness (i.e., inbreeding depression, Keller & Waller, 2002; Crnokrak & Roff, 1999; Charlesworth & Willis, 2009). This can threaten population persistence (Frankham et al., 2002). Genetic diversity is also often correlated with effective population size (N_e_), i.e., the number of individuals that contribute to future generations (Wright, 1931). N_e_ is typically smaller than the census population size due to, e.g., individual variation in reproductive success, arising from unequal sex ratios, mating system and social structure (Palstra & Ruzzante, 2008; Frankham et al., 2011). Low N_e_ increases rate of genetic drift (i.e., changes in allele frequencies due to chance) and inbreeding, which, in turn, accelerates the loss of genetic diversity and the impact of inbreeding depression.

Understanding the relationships between population size and isolation and genetic diversity, effective population size, and inbreeding is thus essential for developing effective conservation strategies, particularly for small and isolated populations (Frankham et al., 2019). Habitat fragmentation reduces populations connectivity and gene flow, which increases the negative impact of genetic drift and inbreeding, with reduced adaptive potential and increased extinction risk as a consequence (Hanski & Gilpin, 1997). Norwegian wild reindeer (*Rangifer tarandus*) are now distributed in 24 more or less isolated populations of varying size, representing a text-book example of the demographic and genetic challenges related to human-caused fragmentation. Historically, these populations were to a larger degree part of a continuous range (Skogland 1994). Before industrial times, harvest reduced population sizes and caused local extirpations, with population genetic implications (Røed et al., 2014). This was followed by habitat loss and fragmentation due to human activities such as infrastructure development and recreational use over the past century (e.g., Nellemann et al., 2001). Such fragmentation is known to disrupt gene flow and increases the risk of genetic drift and inbreeding in ungulates, which in turn can lead to loss of genetic diversity and decreased population viability (Epps et al., 2005; Luikart et al., 2008, Bozzuto et al., 2019).

Barriers such as roads, railways and other infrastructures (Panzacchi et al., 2013) have likely led to genetic isolation and further diversification of the reindeer populations (Kvie et al., 2019). For the smaller populations in particular, this may cause reduced genetic diversity and increased inbreeding, in turn leading to inbreeding depression and reduced adaptive potential (Harrison & Hastings, 1996; Keller et al., 2001). Notably, the increasing demographic and genetic isolation of reindeer populations has occurred in parallel with another anthropogenic disturbance, related to historical and current domestication processes. The 24 present-day populations of ‘wild reindeer’ have contrasting origins (Røed et al., 2014) that can be classified into (original) wild reindeer, semi-domesticated reindeer (released or translocated), and a mixed origin) where semi-domesticated reindeer have intentionally or incidentally been mixed with adjacent wild reindeer populations (Kvie et al., 2019. Such release or mixing of feral individuals into populations of wild genetic origin may have serious negative genetic consequences from a conservation biology perspective (Laikre et al., 2010, Mysterud et al., 2024).

These genetic issues, including the increasing fragmentation into small populations, have so far not received much attention in wild reindeer management and conservation in Norway. In the Norwegian red list for species, the wild reindeer became listed as near threatened in 2021 (Artsdatabanken 2021). This was based solely on demographic changes, i.e., an estimated long-term reduction in total population size that was mainly due to culling and increased hunting quotas related to the detection of chronic wasting disease (CWD). CWD is a lethal prion disease that so far has been detected in two populations, one of which was immediately eradicated by culling organized by the authorities (Mysterud & Rolandsen 2018). With several of the ‘wild reindeer’ populations reaching population sizes as low as 50-100 individuals or even lower, and probably with no or very low gene flow into these populations (Kvie et al., 2019), one may argue for increased conservation focus on maintaining their genetic diversity and facilitating gene flow. However, considering the risk of both further spread of CWD (Mysterud et al., 2020a) and domestic introgression of genetically wild reindeer populations, solving this conservation challenge is not straightforward.

Nevertheless, monitoring and properly managing small and isolated wildlife populations requires state-of-the-art genetic tools to continuously assess levels of genetic diversity, effective population sizes, and inbreeding. Quantifying and understanding these parameters and how they vary within and across populations can inform management strategies aimed at enhancing connectivity and reducing negative effects of habitat loss and fragmentation (Frankham et al., 2019). We developed a custom 625k single nucleotide polymorphism (SNP) array and analyzed 510 individuals across thirteen Norwegian wild reindeer populations, sampled by hunters. For most populations, samples were from two different time periods, ranging from three to 15 years apart. We assessed (loss of) genetic diversity, effective population sizes, and the extent of/changes in inbreeding within each population, and related these linked parameters to population sizes. The use of our high-density SNP array allowed for a detailed and comprehensive analysis of genetic variation across the genome, providing robust estimates of genetic diversity, N_e_, and inbreeding coefficients.

## Methods

### Generating a custom Axiom 625k SNP array

Selection of variants for the SNP array was based on shotgun sequencing data from an unpublished reference panel of 52 reindeer nuclear genomes from diverse Eurasian populations (Bieker et al., in prep.) as well as 100 previously published Svalbard reindeer (*Rangifer tarandus platyrhynchus*) nuclear genomes (Burnett et al. 2023). The sequence data were processed in angsd v0.932 (Korneliussen et al., 2014) and mapped against a caribou reference genome assembly (Taylor et al. 2019). In order to generate a list of variant positions, genotype likelihoods were estimated for the Eurasian and Svalbard reindeer separately with angsd (options *-doGlf 2, -doMajorMinor 1, -SNP_pval 1e-6, -doMaf 1, -doGeno 32, -doPost 1, -GL 2, -remove_bads 1, -uniqueOnly 1, -minMapQ 30, -minQ 20*) and requiring a site to be covered in at least half the individuals. Mappable regions of the reference genome were identified by applying the Genome Analysis Toolkit (GATK) v3.7 (McKenna et al. 2010) CallableLoci function to a merged BAM file consisting of 10 individual BAM files with approximately equal nuclear genome sequencing depth (∼3x) and the options *-mmq 25, -mbq 20, - minDepth 12, -maxDepth 74*. Using this information, variants were excluded from further analysis if they were discovered in reference genome regions where we observed excessive or deficient coverage (higher than 2 times or less than one third of the mean coverage) from the mean sequencing depth across the merged genomic data from 10 individuals in the reference panel, or where alignments generated poor mapping quality (*POOR_MAPPING_QUALITY > 0.1*).

Next, data from a panel of 10 deep-sequenced Svalbard reindeer genomes (Burnett et al., 2023) were used to identify the locations of insertions/deletions (INDELs) in the Svalbard population. To generate a GVCF file for each deep-sequenced sample, GATK v4.1.8.1 HaplotypeCaller was used specifying a minimum mapping quality of 30, option *-stand-call-conf 30*, and a ploidy of 2. Afterwards, a genome database was generated with GATK GenomicsDBImport, followed by a joined SNP call with GATK GenotypeGVCFs using option *-stand-call-conf 30*. Thus, a master list of unique SNP and INDEL positions was constructed. This list was supplemented with a subset of genomic positions from a published 67k SNP array for caribou (Carrier et al., 2022), adding 57,811 polymorphic positions that could be unambiguously lifted over to the caribou genome assembly. Then a custom Python script was used to randomly prune SNPs so that no SNP remained within a 20 bp proximity to another SNP or to a Svalbard panel INDEL, with the pruning algorithm including SNPs according to the following priority criteria (priority ranked from highest to lowest): (1) Svalbard panel SNPs within coding regions, (2) SNPs from the filtered caribou SNP array, (3) Svalbard panel SNPs outside repetitive genomic regions (from genome annotation at www.caribougenome.ca/resources), (4) Eurasian panel SNPs within coding regions, (5) Eurasian panel SNPs outside repetitive genomic regions, and (6) Svalbard panel SNPs within repetitive genomic regions. This pruning process produced a list of 1,995,783 candidate SNPs. Based on the 35bp sequence information on either side of each candidate SNP and the reference caribou genome, the ThermoFisher bioinformatics team classified these candidate SNPs as either “recommended”, “neutral” or “not recommended”, and gave the expected probability of conversion on the forward and reverse strand for each SNP. The SNPs on our array were selected among the ones classified as “recommended” (SNP priority criteria 1-6), “neutral” (SNP priority criteria 1 and 2), or “not recommended” (SNP priority criterion 2). Next, after retaining all remaining priority 1-5 SNPs and the first and last SNP on each scaffold, additional priority (6) SNPs were selected if they were >600 bp away from the two closest SNPs of any other SNP regardless of priority criterion. This resulted in a final selection of 625,857 SNPs (Supplementary Material S1). The mean and median conversion probability for these SNPs were 0.675 and 0.684, respectively, and only 5.8% of the SNPs had a conversion probability below 0.6. The SNPs are separated by a mean of 3,513 bp (median 1,968 bp), with inter-SNP distances ranging from a minimum of 21 bp to a maximum of 100,481 bp. The number of selected SNPs on our custom array of each priority criterion were: 1) 53,157, 2) 39,541, 3) 85,476, 4) 95,912, 5) 37,692, and 6) 314,079. Thus, by including SNPs identified in reindeer across several populations in Eurasia and Svalbard as well as caribou, we developed a custom SNP array with low ascertainment bias suitable for SNP-genotyping in populations across the whole distribution of reindeer globally.

### Study system and populations

Our 13 target populations of ‘wild reindeer’ are ‘wild’ as defined by national management authorities, i.e., reindeer populations that are under hunting regulation and not currently managed as semi-domesticated herds (Table 1, Figure 1). These populations are of different genetic origin, spanning from completely feral/semi-domesticated to a mix of feral and wild mountain reindeer (Kvie et al. 2019). The populations also differ substantially in size, spanning from <50 to thousands of individuals (Table 1). In total n = 510 samples (i.e., unique individuals) were genotyped (Supplementary Material S2). Note that the only information source for ‘population size’ in this study are the ‘population aims’, i.e., the local management’s goal for average winter population size over time, typically based on an approximate assessment of the actual population size. With the considerable variation in population sizes, we believe these estimates are sufficiently accurate for the purpose of the current study.

**Figure 1.**
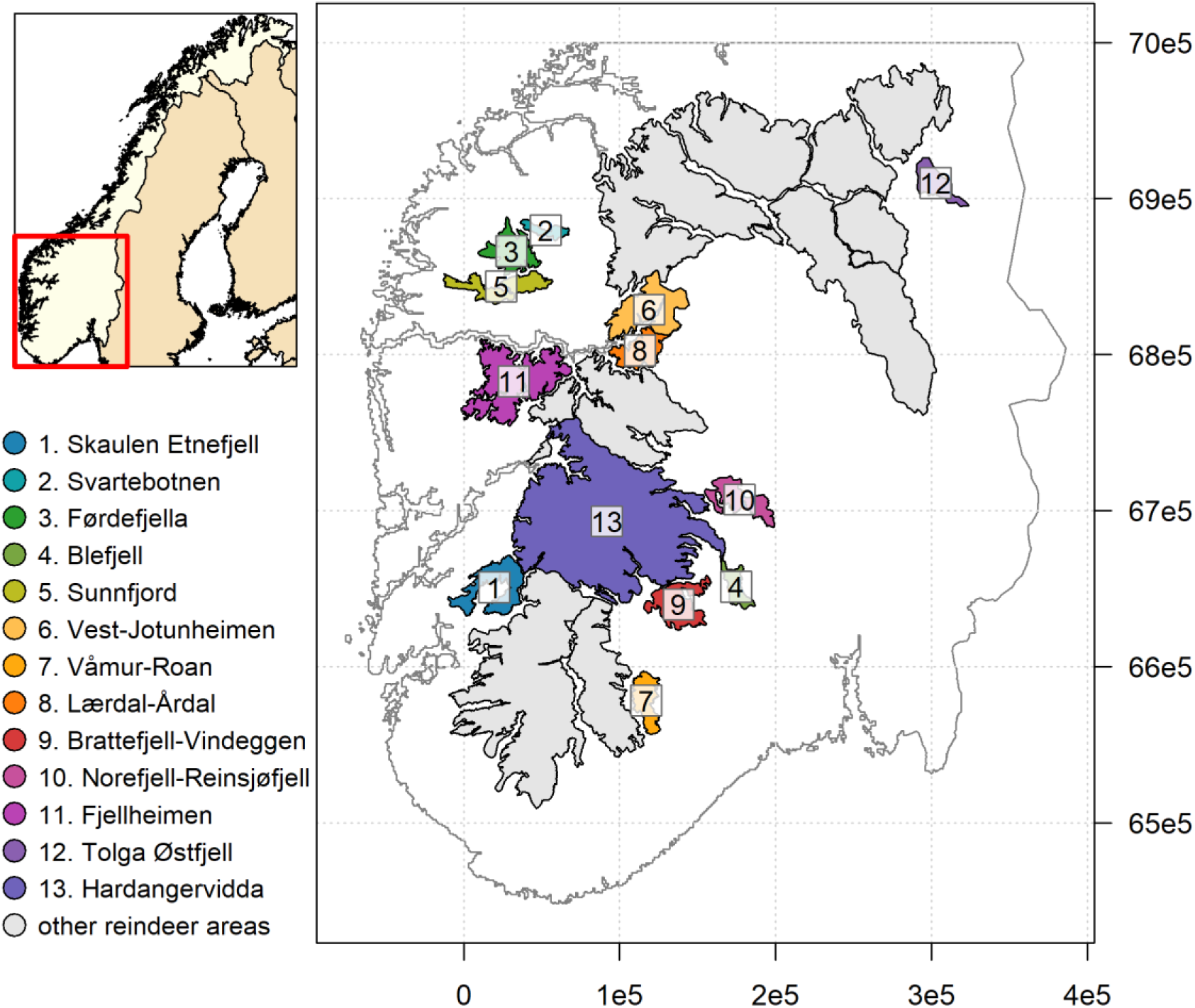
Map showing the distribution and geographical delimitations of the thirteen wild reindeer study populations in Southern Norway that were analyzed in this study. The study populations are ordered from smallest to largest based on approximate population size. X and Y coordinates are projected in UTM zone 33N.

**Table 1.** Sampling information and genetic metrics for 13 reindeer populations in Norway. H_o_ (observed heterozygosity) was estimated using the 107,241 polymorphic SNPs that were mutual across genotyping batches after quality control. Known or assumed genetic origin is divided into D = domesticated or M = mixed wild and domesticated (Kvie et al., 2019).

| Population | Origin | Pop. size (N) | Period | Sample size (n) | Years | $H_o$ (SD) | $N_e$ (95% CI) | $F_{ROH > 8Mbp}$ (95% CI) |
| --- | --- | --- | --- | --- | --- | --- | --- | --- |
| Blefjell | M | 150 | first | 21 | 2003 | 0.310 (0.023) | 8 (3-22) | 0.083 (0.061-0.111) |
|  |  |  | second | 20 | 2017-2019 | 0.301 (0.021) | 25 (14-72) | 0.077 (0.056-0.104) |
| Brattefjell-Vindeggen | M | 550 | first | 28 | 2012-2013 | 0.334 (0.016) | 76 (46-179) | 0.025 (0.015-0.039) |
|  |  |  | second | 33 | 2019 | 0.334 (0.013) | 145 (92-308) | 0.024 (0.015-0.037) |
| Fjellheimen | D | 600 | first | 32 | 2016 | 0.287 (0.010) | 79 (48-177) | 0.024 (0.014-0.036) |
|  |  |  | second | 33 | 2019 | 0.277 (0.020) | 39 (28-61) | 0.047 (0.033-0.064) |
| Førdefjella | D | 100 | first | 9 | 2014 | 0.300 (0.019) | 11 (5-28) | 0.036 (0.016-0.068) |
|  |  |  | second | 13 | 2017-2019 | 0.284 (0.019) | 43 (28-82) | 0.051 (0.029-0.080) |
| Hardangervidda | M | 12000 | first | 0 | - | - | - | - |
|  |  |  | second | 18 | 2021 | 0.351 (0.002) | 2963 (1195-Inf) | 0.001 (0.000-0.008) |
| Lærdal-Årdal | M | 500 | first | 0 | - | - | - | - |
|  |  |  | second | 25 | 2016-2019 | 0.340 (0.013) | 81 (49-200) | 0.016 (0.008-0.028) |
| Norefjell-Reinsjøfjell | D | 570 | first | 26 | 2007 | 0.287 (0.009) | 168 (124-259) | 0.037 (0.024-0.054) |
|  |  |  | second | 27 | 2019 | 0.286 (0.012) | 88 (49-315) | 0.037 (0.024-0.054) |
| Skaulen Etnefjell | D | 50 | first | 16 | 2011-2013 | 0.294 (0.028) | 6 (2-25) | 0.081 (0.056-0.112) |
|  |  |  | second | 23 | 2017-2019 | 0.263 (0.031) | 6 (3-10) | 0.137 (0.109-0.168) |
| Sunnfjord | D | 150 | first | 28 | 2010-2014 | 0.291 (0.025) | 18 (12-28) | 0.043 (0.029-0.061) |
|  |  |  | second | 24 | 2017-2019 | 0.287 (0.023) | 18 (9-47) | 0.052 (0.035-0.073) |
| Svartebotnen | D | 60 | first | 8 | 2012 | 0.266 (0.015) | 33 (16-305) | 0.091 (0.055-0.138) |
|  |  |  | second | 14 | 2017-2019 | 0.252 (0.024) | 30 (19-63) | 0.121 (0.088-0.160) |
| Tolga Østfjell | D | 2000 | first | 0 | - | - | - | - |
|  |  |  | second | 19 | 2016-2019 | 0.301 (0.010) | 293 (142-Inf) | 0.005 (0.001-0.016) |
| Vest-Jotunheimen | D | 400 | first | 0 | - | - | - | - |
|  |  |  | second | 31 | 2018-2019 | 0.306 (0.012) | 101 (78-141) | 0.029 (0.018-0.043) |
| Våmur-Roan | D | 200 | first | 32 | 2013 | 0.317 (0.016) | 50 (35-81) | 0.035 (0.023-0.050) |
|  |  |  | second | 30 | 2018-2019 | 0.309 (0.019) | 34 (22-66) | 0.046 (0.032-0.064) |

For each population, available samples (generally skin or tissue) obtained from hunters were, if possible, divided into an ‘early’ (first) and ‘later’ (second) period. Time of collection varied between populations. For many populations, the time between the first and second sampling period was less than one generation (∼ 5-6 years; Burnett et al, 2023). Some populations lacked samples from an ‘early’ period, which prevented analysis of changes in genetics over time (Table 1).

### SNP genotyping

Samples went through DNA extraction and quality/quantity control and were then genotyped at CIGENE (Norwegian University of Life Sciences) following standard protocols for Axiom genotyping and SNP analysis. Genotyping of the samples from mainland Norway on our custom Axiom array resulted in a total of 214,530 polymorphic SNPs of high quality (classified as Poly High Resolution). One reason why only 34% of our SNPs were polymorphic is probably that SNPs included on our array were selected if they were variable across reindeer and caribou populations spanning Eurasia, Svalbard and Canada. Thus, it is not surprising that many loci were monomorphic within Norwegian populations. The SNP genotype data were quality filtered for further genetic analyses in PLINK v.1.9 (Chang et al. 2015). Individuals with >3% missing genotypes were removed, as well as sample replicates with the highest proportion of missing genotypes. SNPs were excluded when having a minor allele frequency (MAF) <0.01 (n = 1,514) or being significantly out of Hardy-Weinberg equilibrium at p<1e-50 (n = 509). Linkage pruning was performed using the parameters window size = 50, step size = 5, r2 = 0.5. After filtration, a total of 147,934 SNPs were retained. Note that the samples from Hardangervidda were not part of the same genotyping batch as the other populations. The SNPs that passed quality control, therefore, deviated between genotyping batches, which required further filtering (remaining) prior to comparative analyses including Hardangervidda (107,241 SNPs remaining).

### Population genetic analyses

Observed heterozygosity (H_o_) was measured as the proportion of heterozygous SNPs per individual using the *R* package *adegenet* (Jombart and Ahmed 2011). These individual measures of H_o_ were used to estimate percentwise changes in average H_o_ at the population level, using a linear regression model with population and period (early or late), and their interaction, as explanatory variables.

Effective population sizes (N_e_) were estimated with 95% confidence intervals based on the jackknife approach implemented in *NeEstimator* v.2.1 (Nomura 2008, Do et al. 2014). To reduce computational workload, we estimated N_e_ based on the same subset of 10,000 randomly selected SNPs for all populations.

Inbreeding coefficients (F) were estimated from runs of homozygosity (RoH) (Kardos et al. 2016) in *PLINK* using 84,316 out of 214,530 SNPs (i.e. without linkage pruning) occurring on >10 Mbp scaffolds (Burnett et al. 2023), having MAF > 0.01, and using the parameters *--homozyg-window-het* 1 *--homozyg-kb* 500 *--homozyg-snp* 50 *--homozyg-window-snp* 50 *--homozyg-gap* 500 *--homozyg-density* 15 *--homozyg-window-missing* 5. RoHs were aggregated in three size classes – short (0.5-1 Mbp), moderate (1-8 Mbp) and long (> 8Mbp) – reflecting shared ancestry approximately 50-100, 6-50 and < 6 generations ago, respectively, given a recombination rate of ∼1 cM/Mbp (Thompson 2013, Kardos et al. 2017). Changes in recent inbreeding coefficients (F_ROH >8Mbp_) from the first to second sampling period were estimated using a generalized linear model with quasibinomial family and logit link function.

## Results

The estimated genetic variation (H_o_), effective population size (N_e_), and level of inbreeding (F_RoH_) varied strongly between populations (Table 1). H_o_ was positively correlated with population size (Figure 2a) and was generally lower in the second compared with the first sampling period. No population showed a sign of increase in H_o_. Loss of H_o_ over time ranged between 0-10% and was higher the smaller the population (Figure 2b, Table 2). The loss of H_o_ from the first to the second period approached zero for the largest populations.

**Figure 2.**
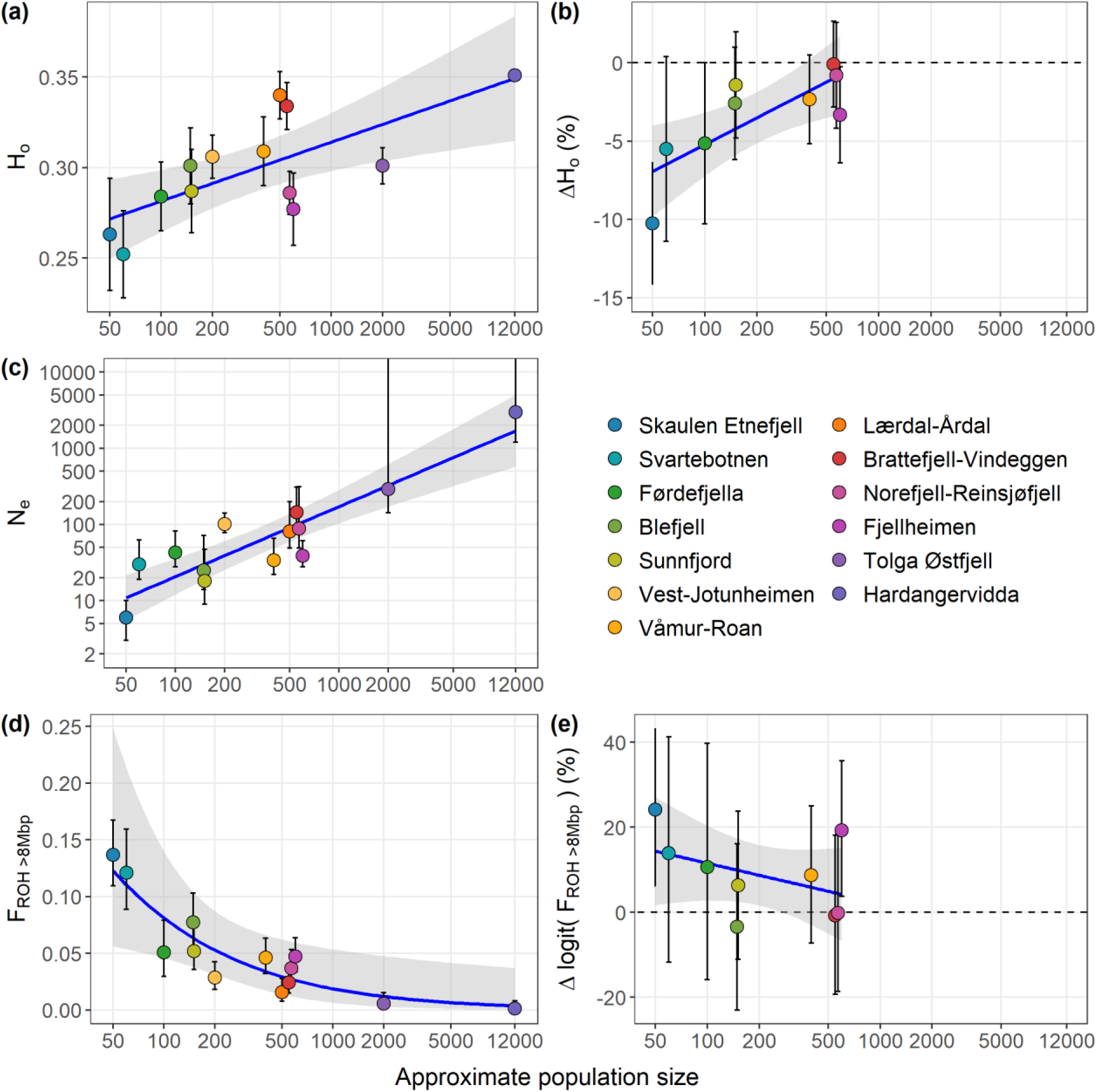
The relationship between approximate actual total population size and (a) genetic diversity (H_o_), (b) change in genetic diversity (percentwise change in H_o_ between the two sampling periods), (c) effective population size (N_e_), (d) recent inbreeding (F_RoH>8Mbp_), and (e) change in recent inbreeding across thirteen wild reindeer populations in Norway. Note that changes in genetic diversity and inbreeding could only be estimated for nine populations. Bars show 95% confidence intervals.

**Table 2.**
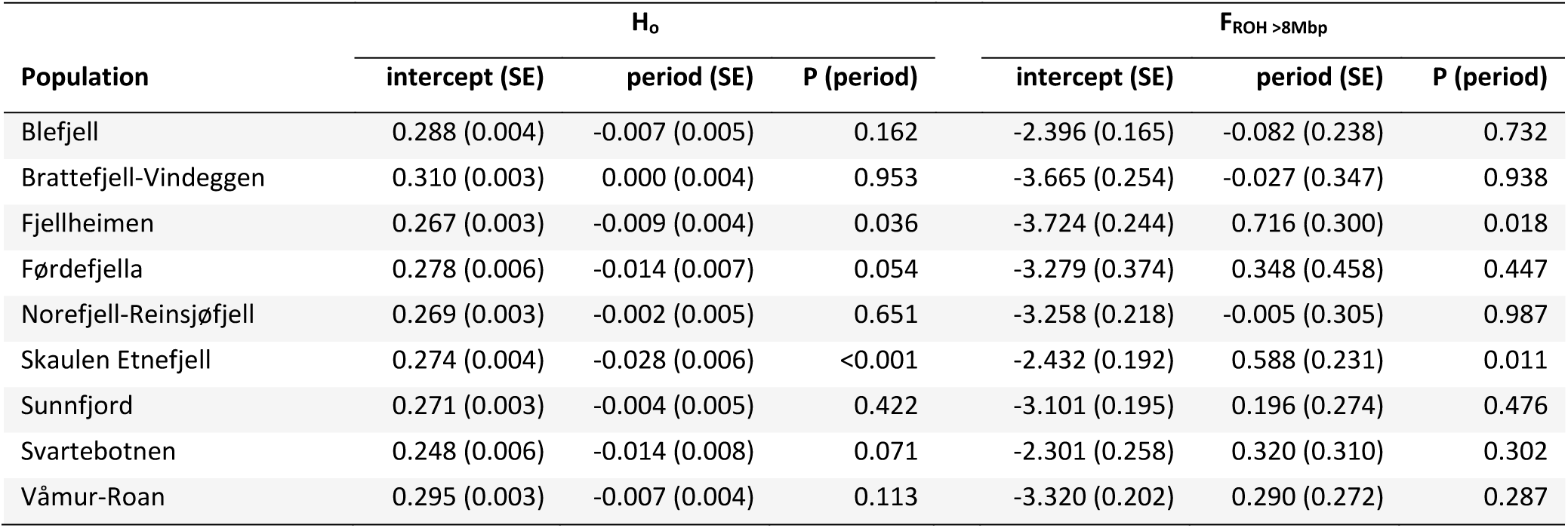
Estimated changes in H_o_ and F_ROH_ (based on RoH>8Mbp) between the first (intercept) and second period in nine reindeer populations in Norway. Estimates for H_o_ are based on individual heterozygosity over 147,934 polymorphic SNPs. Estimates for F_ROH_ are on logit scale.

Mean estimated N_e_ ranged from six to 2,963 individuals (Table 1), with seven out of 13 populations having N_e_<50 in one or both sampling periods. Note, however, that some estimates were highly uncertain, typically (but not only) when based on small sample sizes. As for H_o_, N_e_ was strongly positively related to actual population size (Figure 2c).

The populations in Blefjell, Skaulen-Etnefjell, Sunnfjord, and Svartebotnen had the highest estimated levels of recent inbreeding, measured as F_RoH_ values for RoH>8 Mbp (Table 1, Figure 2d, Figure 3). For most populations, the estimated degree of inbreeding showed some tendency to increase over time (Figure 2e), although a statistically significant change was only found for Fjellheimen and Skaulen-Etnefjell (Table 2). As for N_e_ and H_o_, the variation in estimated recent inbreeding between populations depended on population size, with a higher degree of inbreeding associated with lower population size (Figure 2e). This pattern was less clear for more historical inbreeding (Figure 3).

**Figure 3.**
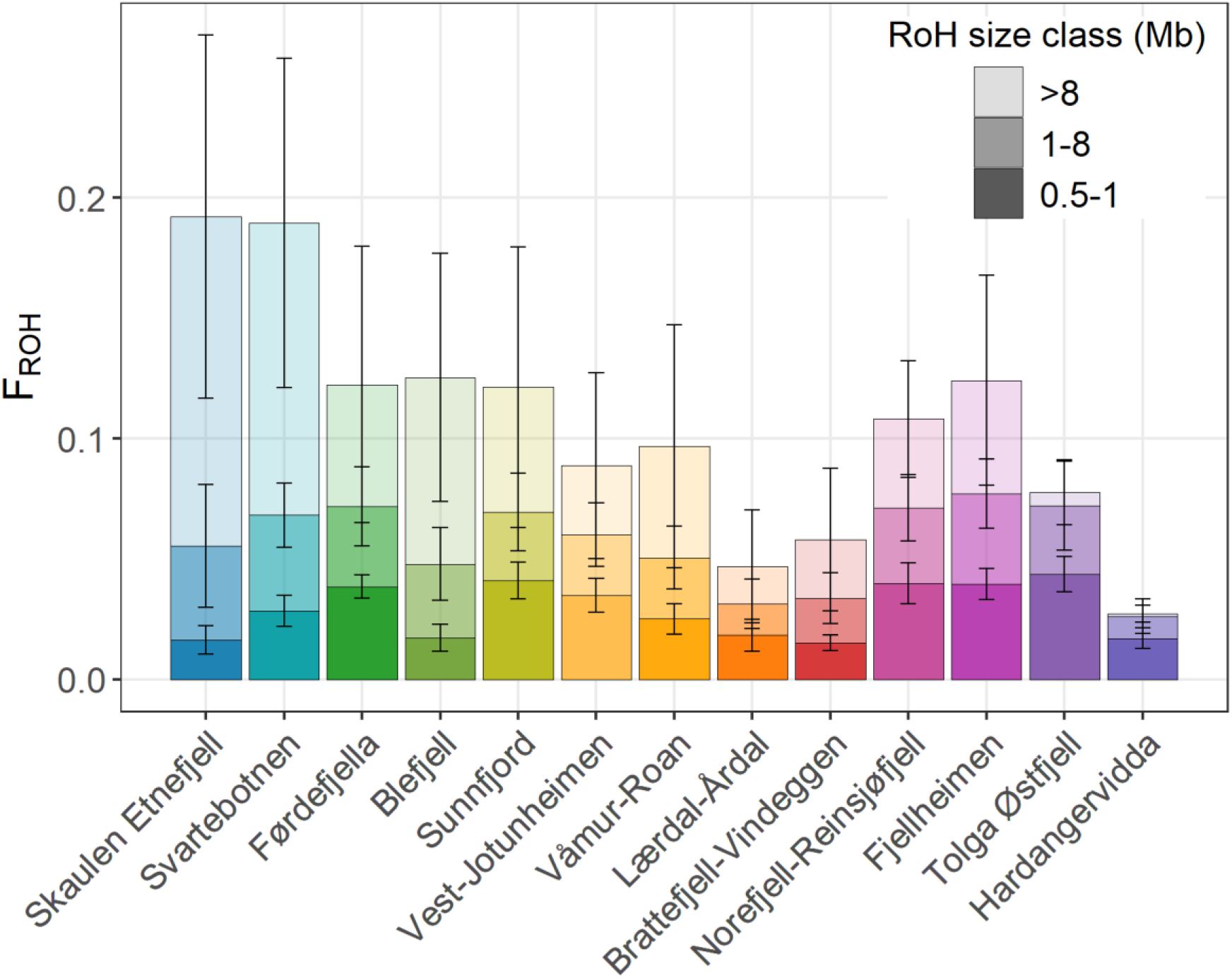
Cumulative inbreeding coefficients F_ROH_ from different Runs of Homozygosity (RoH) size classes (0.5–1 Mbp, 1–8 Mbp and >8 Mbp) across thirteen wild reindeer populations in Norway, sorted by increasing population size. Vertical bars extending from the mean values represent standard deviations.

## Discussion

Habitat loss and fragmentation are increasingly impacting wildlife populations, but the magnitude of effects on the trajectory of genetic diversity has only rarely been assessed comparatively across multiple populations. Here, we designed a custom high-density SNP array for *Rangifer* and genotyped samples from hunted individuals in 13 Norwegian wild reindeer populations to investigate the relationships between population size and genetic diversity, effective population sizes, and inbreeding. The results showed significant variation in these genetic parameters across populations and that population size is a strong determinant of effective population size, inbreeding, and the potential for rapid loss of genetic variation (Tables 1 and 2, Figure 2). Our study provides valuable insights into genetic dynamics, highlighting significant spatiotemporal variation with large implications for conservation strategies aimed at mitigating genetic erosion in such fragmented populations.

The observed correlation between genetic variation and population size underscores the vulnerability of smaller populations to genetic drift that causes loss of genetic diversity. Larger populations consistently exhibited higher H_o_, confirming theoretical expectations and findings from other wildlife studies (Frankham, 1996; Reed & Frankham, 2003). Notably, no population showed an increase in H_o_ over time, with losses ranging from 0-10%, where the non-linear relationship between population size and H_o_ indicates a steeper decline in genetic diversity as population size decreased. An estimated 10% loss of genetic variation (in the Skaulen Etnefjell population) over a period of only approximately one generation is very high (Leigh et al., 2019). This pattern of reduced H_o_ was particularly pronounced in smaller populations, consistent with studies on other species of conservation concern (Johnson et al., 2010; O’Brien et al., 1985). Cautious interpretation is, however, needed for some populations with ongoing sub-fragmentation, because information on the analyzed samples was constrained geographically to their respective total population range, or ‘reindeer area’ (as delimited in Figure 1). Thus, we cannot rule out the possibility that the distribution of sample locations between isolated subareas within the population range differs between the first and second sampling periods, which means that variation in genetic metrics within populations and estimated changes over time could be influenced by spatial effects. The same problem potentially applies where no significant changes were found. Thus, especially the results from Fjellheimen, Skaulen-Etnefjell, and Sunnfjord should be interpreted with caution, as these populations encounter major barriers to movement within their range.

Effective population size estimates also varied widely and – unsurprisingly – according to the population size, emphasizing the critical role of population size in maintaining genetic stability.

Populations with high loss of genetic diversity and very low effective population sizes, such as Skaulen Etnefjell, Svartebotnen, and Førdefjella, are particularly at risk of inbreeding and genetic drift, which can further exacerbate genetic erosion and reduce adaptive potential (Charlesworth & Willis, 2009; Palstra & Ruzzante, 2008). As expected, estimates of recent inbreeding were relatively high in these populations, highlighting their geographic isolation and ongoing genetic erosion. There was an overall tendency for increased inbreeding with time (i.e., between the two sampling periods), and with an overall stronger increase in the smaller populations. Note that, in some of these populations, the time between sampling periods was very short (even less than one generation), further highlighting the apparently rapid loss of genetic variation. High inbreeding coefficients are associated with reduced fitness and increased risk of detrimental impacts of inbreeding depression, as documented in other species (Liberg et al., 2005, Hewett et al., 2024). The correlation between population size and inbreeding levels thus further underscores the importance of maintaining sufficiently large population sizes to minimize inbreeding and its deleterious effects.

Notably, estimates of more historical inbreeding (F_RoH_ for RoH size classes 0.5-1 and 1-8 Mbp) followed a different pattern across populations (Figure 3), i.e., not that strongly related to present-day population size. This may appear to be related to genetic origin. The mixed-origin populations Blefjell, Brattefjell-Vindeggen, Hardangervidda, and Lærdal-Årdal (see Table 1) had low F_RoH_ values for RoH = 0.5-1 (ca. 50-100 generations ago) in particular, when compared to those with the semi-domesticated origins. This pattern could be related to the selection processes and loss of genetic variation from domestication a few hundred years ago (e.g., Røed et al., 2014), but more in-depth analysis of population-genetic structuring and the genetic origins of these populations are needed to support this.

The spatiotemporal variation in genetic diversity, effective population size, and inbreeding documented here indicates a need for targeted conservation actions to preserve these populations. Regularly monitoring genetic parameters to assess the effectiveness of conservation interventions and subsequently to adapt strategies as needed can help detect further signs of genetic erosion and inbreeding depression, enabling timely management actions. Management could also implement genetic rescue by introducing individuals from other populations to increase genetic diversity and N_e_ and reduce inbreeding levels. This strategy has been successful in other conservation contexts, such as the genetic restoration of the Florida panther (Johnson et al., 2010). Furthermore, enhancing habitat connectivity by establishing wildlife corridors and mitigating barriers such as roads and fences could facilitate gene flow between populations. This approach is often crucial for maintaining genetic diversity and reducing inbreeding (Hanski & Gilpin, 1997; Crooks & Sanjayan, 2006).

However, planning and implementing management actions involving gene flow among reindeer populations requires cautious consideration. Most of the studied populations, including the smaller ones with the largest demographic and genetic challenges, descend from semi-domesticated reindeer that previously were either introduced (once, or on several occasions) or released into the wild. Earlier mixing of semi-domesticated reindeer or reindeer with semi-domesticated origin into populations of genetically wild mountain reindeer has led to domestic genetic introgression (yet also increased genetic variation), including in the largest population, Hardangervidda. Most populations studied here share ranges adjacent to Hardangervidda and/or other populations with mixed origin, separated by human infrastructure such as roads. Thus, even if the intention of reducing barriers would be to increase gene flow only in one direction, management actions increasing dispersal opportunities could also contribute to further domestic introgression of populations that have previously experienced only limited gene flow from semi-domestic herds. This would lead to a situation similar to the potential use of semi-domesticated animals in reintroduction programs (Mysterud et al., 2024). In addition, Hardangervidda is one of the two reindeer populations in Norway where CWD has been detected. Restricting dispersal of individuals to other populations is currently a focal management goal, along with intensified male-biased harvesting, resulting in a skewed sex ratio, reduced population size, and possibly reduced genetic diversity in the long run (Mysterud et al., 2020a, 2020b, Kvalnes et al., 2024).

Our study has highlighted the substantial genetic challenges faced by several Norwegian reindeer populations due to a combination of their small population size and lack of gene flow. It also underscores some of the conservation dilemma characterizing this system, as management actions to reduce barriers between populations (with intentions to increase gene flow and genetic diversity) currently contradicts other major conservation goals, such as mitigating the spread of CWD and avoiding domestic introgression into wild-origin populations. Thus, maintaining genetic diversity and connectivity can be urgent in a strict population-level context, while at the species/national level, other management priorities may overrule this. Nevertheless, our study provides an example of the applicability of cost-efficient, high-density SNP arrays as a tool to monitor and detect genetic conservation challenges in wildlife populations.

## Acknowledgments

The development of the 625k SNP array was financed by the Research Council of Norway (RCN, grants 223257, 276080, 302619, 325589, and 343398), the Centre for Biodiversity Dynamics and the Gjærevoll Centre for Biodiversity Foresight Analysis at the Norwegian University of Science and Technology (NTNU), and the Norwegian Institute for Nature Research (NINA). The DNA extraction and genotyping of the reindeer in this study, as well as the data analysis and writing of this paper, were financed by the Norwegian Environment Agency, NINA, and RCN (grants 343398 and 325589).

## Data availability

The Axiom 625k SNP array for *Rangifer tarandus* (Supplementary Material S1, https://doi.org/10.5061/dryad.v6wwpzh47) and SNP genotype data for the 510 reindeer used in this study (Supplementary Material S2, https://doi.org/10.5061/dryad.pzgmsbcwj) will be published on Dryad at the time of publication in a journal and are, until then, available upon request.

## Supplementary Material

**Supplementary Material S1.** Hansen BB et al. (Fourthcoming 2024). *Rangifer tarandus* Axiom 625k SNP array [Dataset]. Dryad. https://doi.org/10.5061/dryad.v6wwpzh47.

**Supplementary Material S2.** Hansen BB et al. (Fourthcoming 2024). SNP genotype data to “Rapid loss of genetic variation and increased inbreeding in small and isolated populations of Norwegian wild reindeer” [Dataset]. Dryad. https://doi.org/10.5061/dryad.pzgmsbcwj.

